# Single-Cell RNA Sequencing of Batch Chlamydomonas Cultures Reveals Heterogeneity in their Diurnal Cycle Phase

**DOI:** 10.1101/2020.09.15.298844

**Authors:** Feiyang Ma, Patrice A. Salomé, Sabeeha S. Merchant, Matteo Pellegrini

## Abstract

The photosynthetic unicellular alga Chlamydomonas (*Chlamydomonas reinhardtii*) is a versatile reference for algal biology because of the facility with which it can be cultured in the laboratory. Genomic and systems biology approaches have previously been used to describe how the transcriptome responds to environmental changes, but this analysis has been limited to bulk data, representing the average behavior from pools of cells. Here, we apply single-cell RNA sequencing (scRNA-seq) to probe the heterogeneity of Chlamydomonas cell populations under three environments and in two genotypes differing in the presence of a cell wall. First, we determined that RNA can be extracted from single algal cells with or without a cell wall, offering the possibility to sample algae communities in the wild. Second, scRNA-seq successfully separated single cells into non-overlapping cell clusters according to their growth conditions. Cells exposed to iron or nitrogen deficiency were easily distinguished despite a shared tendency to arrest cell division to economize resources. Notably, these groups of cells recapitulated known patterns observed with bulk RNA-seq, but also revealed their inherent heterogeneity. A substantial source of variation between cells originated from their endogenous diurnal phase, although cultures were grown in constant light. We exploited this result to show that circadian iron responses may be conserved from algae to land plants. We propose that bulk RNA-seq data represent an average of varied cell states that hides underappreciated heterogeneity.

**One-sentence summary:** We show that single-cell RNA-seq (scRNA-seq) can be applied to Chlamydomonas cultures to reveal the that heterogenity in bulk cultures is largely driven by diurnal cycle phases

The author responsible for distribution of materials integral to the findings presented in this article in accordance with the policy described in the Instructions for Authors (www.plantcell.org) is: Matteo Pellegrini

## INTRODUCTION

Transcriptome analysis in the green unicellular alga Chlamydomonas (*Chlamydomonas reinhardtii*) has proliferated since the genome was released in 2007 (Merchant et al., 2007). Since then, dozens of experiments have been conducted that aimed to describe the changes in gene expression in response to changes in nutrient availability such as nitrogen (Plumley and Schmidt, 1989; Blaby et al., 2013; Park et al., 2015), sulfur (González-Ballester et al., 2010), phosphorus (Bajhaiya et al., 2016), acetate (Bogaert et al., 2019) and essential metals (Urzica et al., 2012; Blaby-Haas et al., 2016; Merchant et al., 2006; Malasarn et al., 2013; Blaby-Haas and Merchant, 2012; Kropat et al., 2015), as well as changes that occur in response to light (Tilbrook et al., 2016) or across the diurnal cycle (Zones et al., 2015; Strenkert et al., 2019) and following chemical treatments (Blaby et al., 2015; Ma et al., 2020; Wittkopp et al., 2017). A common feature of the prior studies is the use of bulk RNA-seq obtained from the sequencing of RNA extracted from pools of cells, which is due to the technical necessity to meet the material requirements for library preparation. Changes in transcript levels therefore reflect the average behavior of the culture and may not accurately inform on the extent of cell-to-cell variation that might exist in these samples.

The recent development of single-cell RNA sequencing techniques (scRNA-seq) has gained in popularity to counter the innate limitations of bulk RNA-seq. In Arabidopsis (*Arabidopsis thaliana*) and yeast (*Saccharomyces cerevisiae),* the comparison of bulk RNA-seq and scRNA-seq results has highlighted the heterogeneity of cell populations. For instance, the characterization of yeast culture responses to stress has uncovered variability in gene expression between cells, which may shape how well they cope with the introduced stressor (Gasch et al., 2017). Individual yeast cells also do not age evenly within cultures, again highlighting the heterogeneity of bulk cultures (Zhang et al., 2020). Likewise, in Arabidopsis, the profiling of single root cells has revealed the stochasticity reflecting their developmental trajectories, although each cell type can be efficiently identified by comparing scRNA-seq and bulk RNA-seq data (Zhang et al., 2019; Shulse et al., 2019). In both Arabidopsis and yeast, the cells under investigation are surrounded by a physical barrier that needs to be removed prior to RNA extraction and library construction. In the case of Arabidopsis, the cell wall is digested by a mixture of enzymes for 60 min; protoplast isolation ahead of scRNA-seq may therefore introduce variation in the gene expression profile of single cells that needs to be taken into account during subsequent analysis, especially for short-lived RNAs.

As a photosynthetic unicellular alga, Chlamydomonas presents an ideal system for the application of single-cell RNA sequencing (scRNA-seq) to discover whether cultures exhibit similar stochasticity in their transcriptome as Arabidopsis root cells, yeast, or mammalian cells. Although the alga can be easily synchronized to a 24 h cell division cycle by growth under light-dark cycles (12 h light/12 h dark), the vast majority of experimental conditions rely on cells grown in constant conditions. In addition, cultures need to be refreshed often so as to keep cells in an actively growing state. It is assumed that such cultures are globally asynchronous and represent a mixture of cells in various phases along the diurnal and cell cycles. However, this assumption has never been tested emprically.

We describe here the scRNA-seq analysis of gene expression for almost 60,000 cells derived from three growth conditions and two Chlamydomonas strains. We report that scRNA-seq successfully captures the same gene expression signatures as bulk RNA-seq approaches. We further show that cells experiencing distinct growth conditions cluster independently from one another. Finally, we determine that bulk Chlamydomonas cultures grown in constant light are far from homogeneous and exhibit instead substantial variation in their diurnal cycle, although the distribution of these phases is not uniform. We then use the preferential diurnal phase exhibited by cells to demonstrate the likely conservation of circadian iron responses in Chlamydomonas, as diurnal phases are globally lagging in iron deficient algal cells, as seen in Arabidopsis (Salomé et al., 2013; Hong et al., 2013; Chen et al., 2013).

## RESULTS

### Single-Cell RNA Sequencing (scRNA-seq) of Chlamydomonas Cells Reflects their Iron Nutritional

To determine whether single-cell RNA sequencing (scRNA-seq) methodology is applicable off-the-shelf for profiling Chlamydomonas cultures, we tested the cell wall-deficient strain CC-5390 under two contrasting conditions: iron-replete (Fe+), and iron deficient (Fe–). We grew a single culture for 3 d in constant light and in Fe+ conditions before splitting the culture into Fe+ and Fe– cultures. We measured cell density after 23 h and adjusted it to 1,200 cells/mL for Gel Bead in Emulsion (GEMs) formation and single-cell library preparation. We reasoned that 1 d in the complete absence of Fe would be sufficient to induce a strong Fe deficiency response (Urzica et al., 2012), but would not be as drastic as prolonged Fe deficiency from the time of initial inoculation. To test reproducibility, we also generated a third sample consisting of a mixture of the two samples at equal cell density, and proceeded with GEMs alongside the Fe+ and Fe– samples.

After sequencing and mapping reads to the Chlamydomonas reference genome (version v5.5), we counted 28,690 cells across the three samples, from which we detected an average of 823 genes and 3,344 unique molecular identifiers (UMIs) per cell (Figure 1A). The contribution of mitochondrial and chloroplast transcripts to UMIs was low (0.23% for mitochondria and 0.91% for chloroplasts, Figure 1A), consistent with the initiation of reverse transcription from an oligo(dT) primer (Gallaher et al., 2018).

**Figure 1.**
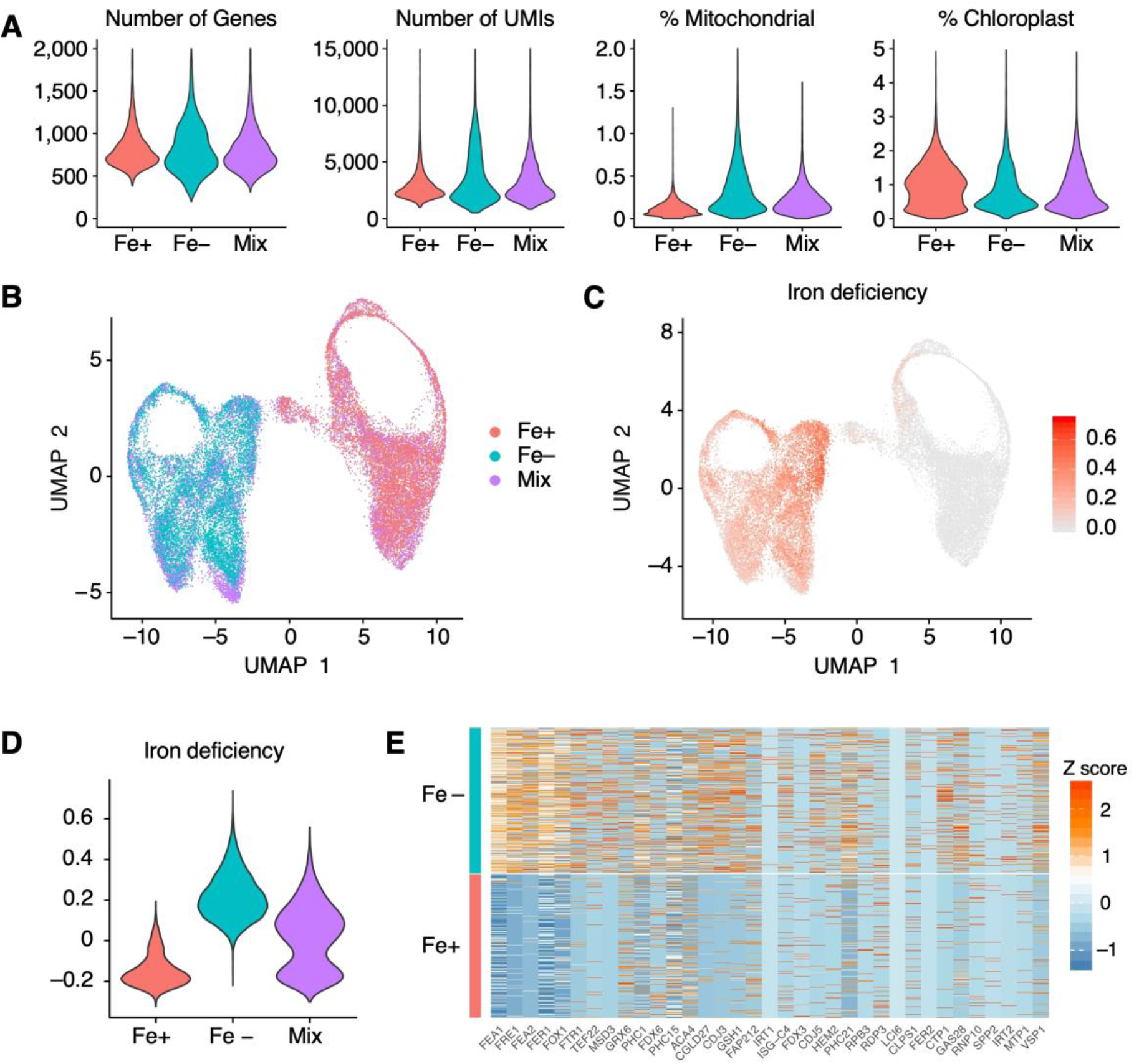
Single-Cell RNA Sequencing Properly Separates Chlamydomonas Cells According to their Iron Nutritional Status. In a first experiment, we grew Chlamydomonas strain CC-5390 in Fe-replete (Fe+) conditions before being transferred to Fe+ or Fe-limited conditions (Fe–) for 23 h. Cells were then processed for scRNA-seq, starting with Gel in Bead Emulsion (GEMs) formation in the 10X Genomics pipeline. **(A)** Characteristics of sequencing results from Chromium Single Cell 3’ gene expression libraries (first experiment). Violin plots report the number of genes, number of unique molecular identifiers (UMIs), and the percentage of gene expression estimates coming from the mitochondrial and chloroplast organelles in Fe+ (pink), Fe– (teal) and an equal mix of cells from Fe+ and Fe– cultures (Mix, purple). **(B)** Uniform manifold approximation and projection (UMAP) plot for the 28,690 sequenced cells, colored by sample: Fe+, pink; Fe–, teal; Mix: purple. Each dot represents one cell. **(C)** UMAP plot of the iron deficiency module score, which includes genes highly induced by Fe deficiency (Urzica et al., 2012). Dark red indicates individual cells with a high iron deficiency module score and thus in a Fe-limited nutritional state. **(D)** Iron deficiency module score for each sample, shown as violin plots. Fe+, pink; Fe–, teal; Mix: purple. Note the bimodal distribution of the Mix sample. **(E)** Heatmap representation of normalized gene expression estimates for of genes induced under Fe deficiency in Fe+ and Fe– cells. Each horizontal line indicates the expression of the listed gene in one cell.

Since the scRNA-seq dataset consisted of expression information from 16,982 genes across about 30,000 cells, we next performed a Uniform Manifold Approximation and Projection (UMAP) dimensionality reduction using the R package Seurat (Stuart et al., 2019). Fe+ and Fe– cells formed two clearly separated groups, while the mixed cells sample was equally divided between the first two groups and closely overlapped with them in the UMAP plot (Figure 1B). These results demonstrated that scRNA-seq 1) successfully separated cells according to their nutritional status (Fe replete or Fe deficient), and 2) had very good technical reproducibility between libraries processed in parallel, as evidenced by the overlap between the mixed cells samples and the two test groups.

To validate the observation that scRNA-seq captured the Fe nutritional status of our samples, we calculated an iron deficiency module score (Stuart et al., 2019) for each cell using genes induced under Fe deficiency previously identified using bulk RNA-seq (Urzica et al., 2012). A module score calculates the average expression of a given gene list, subtracted by the aggregated expression of randomly-sampled control genes. We discovered that Fe– cells exhibited a much higher iron deficiency compared to Fe+ cells, supporting the ability of scRNA-seq to capture expression differences resulting from distinct culture conditions (Figure 1C). The mixed cells sample showed a bimodal distribution for the iron deficiency module score, in agreement with the equal contribution of Fe+ and Fe– cells (Figure 1D).

We also plotted the expression of a number of iron-related genes across all cells, shown as a heatmap in Figure 1E. We observed strong induction for genes encoding various components of the Fe assimilation machinery, such as the *Fe ASSIMILATORY (FEA*) genes *FEA1* and *FEA2,* the *FERRIC REDUCTASE FRE1,* the multicopper oxidase *FOX1* and the Fe permease *FE TRANSPORTER (FTR1).* Other highly expressed genes across Fe– cells included *TEF22,* which is divergently-transcribed from the same promoter sequences as *FEA1;* the low-Fe induced *MANGANESE SUPEROXIDE DISMUTASE 3 (MSD3);* the Chloroplast DnaJ-like *CDJ3* and *CONSERVED IN THE GREEN LINEAGE 27 (CGLD27*) (Urzica et al., 2012). Likewise, the *COPPER-TRANSPORTING P-type ATPase CTP1* was highly expressed only in Fe– cells. CTP1 is predicted to load Cu into FOX1 for full Fe-deficiency responses (Merchant et al., 2006; La Fontaine et al., 2002; Eriksson et al., 2004). The high-affinity Fe transporter *IRT1* was seldom expressed in either Fe+ or Fe– cells, although the related transporter gene *IRT2* was induced in a large fraction of Fe– cells (Figure 1E). Finally, we noted high expression of a number of genes encoding cell wall-associated proteins: cell wall pherophorin-C (*PHC*) *PHC1* and *PHC21,* vegetative hydroxyproline-rich *VSP1,* and *GAMETE-SPECIFIC 28 (GAS28*) (Rodriguez et al., 1999; Waffenschmidt et al., 1993); and plasma membrane proteins such as autoinhibited Ca2+-ATPase 4 (*ACA4), METAL TRANSPORT PROTEIN1 (MTP1*) and *LOW CO2-INDUCED 6 (LCI6).* We interpret these highly induced genes as being part of the stress response of a Chlamydomonas strain lacking a cell wall. scRNA-seq therefore efficiently captures comparable changes in the transcriptome as bulk RNA-seq when Chlamydomonas cells are grown in Fe+ and Fe– conditions.

### scRNA-seq Also Recapitulates Nitrogen Deficiency Bulk RNA Sequencing Signatures

In a second independent experiment, we grew CC-5390 cells under replete conditions for both Fe and nitrogen (N), and then split the cultures into Fe and N replete (control), Fe– (with full N supply) and N deficiency (N–, with full Fe supply, as technical duplicates) 23 h before processing cells for GEMs. After sequencing, we counted 19,140 cells across the four samples, from which we detected an average of 4,181 UMIs and 694 genes per cell. UMAP dimensionality reduction identified three clearly separated clusters, corresponding to control cells (Fe+ N+), Fe-deficient cells (Fe–) and N-deficient cells (N-Fe+) (Figure 2A). These results indicated that scRNA-seq consistently produced distinct cell clusters for Fe+ and Fe– cells across multiple experiments (Figure 1B, Figure 2A). In addition, N– cells formed a cluster that did not overlap with either Fe+ or Fe– cells, suggesting a transcriptome signature that is unique to each growth condition. Finally, we again observed good technical reproducibility, as the two replicates for N– cells closely overlapped.

**Figure 2.**
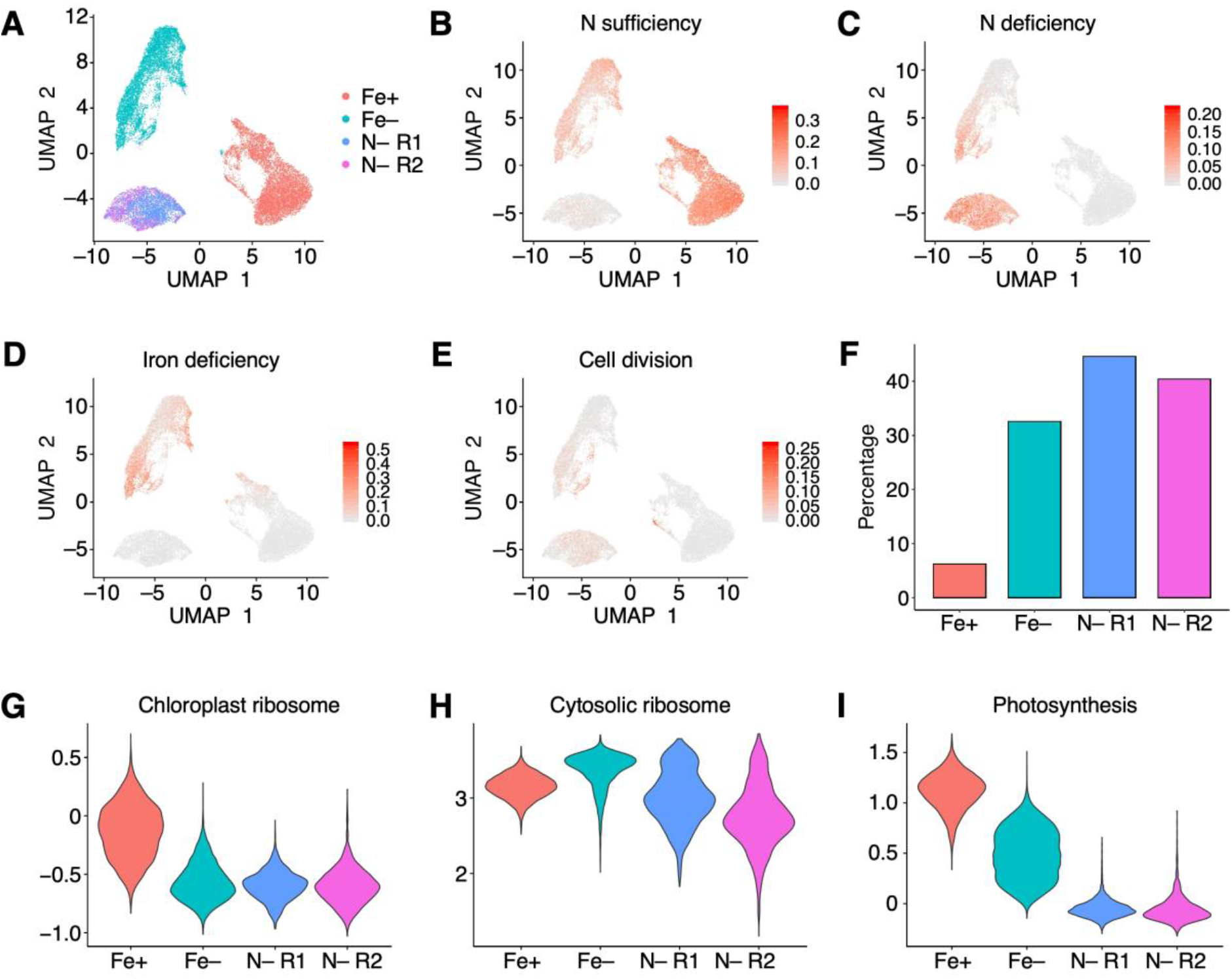
Single cell RNA sequencing Captures Bulk RNA Sequencing Signatures of Nitrogen Deficiency. We grew Chlamydomonas strain CC-5390 in nitrogen (N) and Fe-replete conditions before exposing cells to N deficiency (but full Fe supply) and Fe limitation (with full N supply) for 23 h. **(A)** UMAP plot for 19,140 sequenced cells, colored by sample: Fe+ and N+, red; Fe– and N+, teal; N– and Fe+, purple and magenta (two technical replicates: N– R1 and N– R2). **(B)** UMAP plot of the N sufficiency module score, which includes genes strongly repressed by N deficiency and/or induced by N sufficiency (Schmollinger et al., 2014). Dark red indicates individual cells with a high N sufficiency module score and are thus N– replete. **(C)** UMAP plot of the N deficiency module score, which includes genes highly induced by N deficiency (Schmollinger et al., 2014). Dark red indicates individual cells with a high N deficiency module score and thus in a N-limited nutritional state. **(D)** UMAP plot showing of iron deficiency module score, using the same gene list as in Figure 1. **(E)** UMAP plot showing of cell division module score, based on a list of genes involved in DNA replication and chromosome segregation with a mean diurnal phase of 12-14 h (using dawn as time 0). **(F)** Percentage of cells with a high cell division score across the Fe+, Fe– and N– samples. We included cells with a positive cell division module score. **(G-I)**Module score across all samples for chloroplast ribosome protein genes (*RPGs*) (**G**), cytosolic ribosomes (**H**) and photosynthesis-related genes (**I**). The chloroplast and cytosolic *RPG* module score includes all nucleus-encoded plastid-localized or cytosolic RPG subunits, respectively. The photosynthesis module score is derived from all nucleus-encoded photosystem I and II components, as well as chlorophyll biosynthetic genes and M factors.

To investigate whether scRNA-seq accurately captured the behavior of N status signature genes identified by bulk RNA-seq, we calculated module scores using two gene lists: genes repressed under N deficiency (and thus induced under N sufficiency conditions; Figure 2B) and genes induced under N deficiency (Figure 2C). Both Fe+ and Fe– cells showed a high N sufficiency module score, although Fe+ cells appeared to exhibit a higher score than Fe– cells (Figure 2B). In agreement, a subset of Fe– cells did display a significant module score for N deficiency genes, as expected due to the rearrangement of the photosynthetic apparatus in response to Fe deficiency (Moseley et al., 2002). Notably, N– cells were characterized by a very low module score for N sufficiency marker genes and a high module score for N deficiency genes, thus validating their clustering into a group separate from those of control and Fe– cells (Figures 2B, 2C).

The Fe module score was high in Fe– cells, further confirming the UMAP clustering results (Figure 2D). As Fe– and N– cells would be predicted to stop dividing rapidly to maintain their nutritional quotas (Street and Paytan, 2005), we calculated a module score for genes specifically involved in cell division (minichromosome maintenance (MCM) complex, DNA replication, and Structural Maintenance Of Chromosome (SMC)-encoding genes). Overall, few cells showed a cell division signature, but they largely belonged to the Fe– and N– clusters (Figure 2D). We also observed a sub-group of Fe– cells with a strong cell division module score. We hypothesized that these highlighted cells were arrested prior to entry into cell division proper due to Fe or N deficiency. To test this hypothesis, we calculated the percentage of cells with a positive cell division module score for each sample: 30-40% of Fe– and N– cells fulfilled this criterion, consistent with cell cycle arrest to prevent loss of nutrients (Figure 2F). By contrast, only ~7% of Fe+ cells division had a high cell division module score, as expected for an even distribution of cells along the various stages of the cell cycle.

Due of the high abundance of the photosynthetic apparatus, with a stoichiometry of 1 × 10^6^ molecules per cell, photosynthetic proteins constitute a high draw on the amino acid pool and on the Fe pool because of their high Fe content. Therefore, Fe and N deficiency are expected to have a strong negative effect on the synthesis of the photosynthetic apparatus, and especially in the case of N deficiency, the translation apparatus. We therefore calculated module scores for genes of the photosynthesis apparatus, as well as for ribosomal protein genes (*RPGs*). While mitochondrial *RPGs* showed a constant module score across all conditions (Supplemental Figure 1A), chloroplast *RPGs* were associated with a substantially reduced module score under Fe or N deficiency (Figure 2G). These results are consistent with the cellular response to each nutritional deficit: Fe deficiency will limit chloroplast development, while N deficiency will cause a global reallocation of N resources away from N-rich proteins such as ribosomes (Siersma and Chiang, 1971; Martin et al., 1976) or photosynthetic proteins (Plumley and Schmidt, 1989). This latter hypothesis was also reflected in the module score for cytosolic *RPGs,* which was much lower in N– cells relative to N+ cells (Figure 2H). Finally, the module score for photosynthetic genes recapitulated nicely the known physiological state of each group of cells, with Fe+ cells showing a high photosynthesis module score that decreased in Fe– cells (Figure 2I). N– cells experienced an even stronger repression of the photosynthetic apparatus, with a mean module score close to 0 (Figure 2I). These results independently confirmed the module scores calculated for N sufficiency and deficiency, as several genes encoding photosynthetic components (for example, Light Harvesting Complex proteins 7 LHCAs and 4 LHCBs) are included in the N sufficiency list (Moseley et al., 2002; Peltier and Schmidt, 1991).

Although N deficiency is a routinely employed growth condition to induce the production of lipids in Chlamydomonas, we detected no changes in a module score for lipid biosynthetic genes (Supplemental Figure 2B), suggesting that 1 d of growth in the absence of N was not sufficient to promote lipids biosynthesis, although cultures experienced clear signs of N deficiency, as evidenced by severe chlorosis.

Together, these results demonstrate that scRNA-seq can sort individual cells according to their transcriptional profile in response to multiple stresses and that Fe– and N– cells are arrested before the completion of cell division, likely so as not to dilute their limiting resources and/or because they do not have the necessary resources to multiply.

### Diurnal Rhythmic Oscillations Explain Much of the Heterogeneity of Batch-Cultured Cells

One of the primary advantages of scRNA-seq is that it can reveal the heterogeneity between cells, while bulk RNA-seq only captures the average expression across all cells. In both experiments, we observed clear heterogeneity in both Fe+ and Fe– cells. To explore the source of this heterogeneity in more detail, we ran unsupervised clustering on Fe+ cells from the first experiment and obtained 15 clusters (Figure 3A). Notably, many clusters organized around a closed circle, which indicated that cells might occupy different states within a cycle. We observed a similar circle in the second experiment (Figure 2A) and noted that a fraction of cells appeared to be primed for cell division based on the cell division module score (Figure 2E, 2F). We therefore expanded our analysis to cover all possible phases of the diurnal cycle, using diurnal phases from two recent diurnal time-courses (Zones et al., 2015; Strenkert et al., 2019). We calculated the module score for genes within 1 h time bins every other h, from 0 to 24 h, for each of the five clusters along the circle (clusters # 0- #5, Figure 3A). As shown in Figure 3B, the resulting phase module scores followed a clear pattern that ordered the clusters along the diurnal cycle, with cluster #0 exhibiting a phase close to dawn and clusters #2 and #3 showing a phase close to dusk. We also plotted representative module scores in UMAP plots (Figure 3C). Most cells occupied time bins between 4 and 8 h after lights-on. Smaller cell populations had time signatures closer to 14 h after dawn (largely overlapping with cluster #2), 18 h (corresponding to clusters #3 and #4) and 20 h (matching clusters #5 and #0). As expected for cells progressing through a ~ 24 h rhythm, module scores for the phase bins at 0 h and 24 h were very similar in our analysis (Figure 3C).

**Figure 3.**
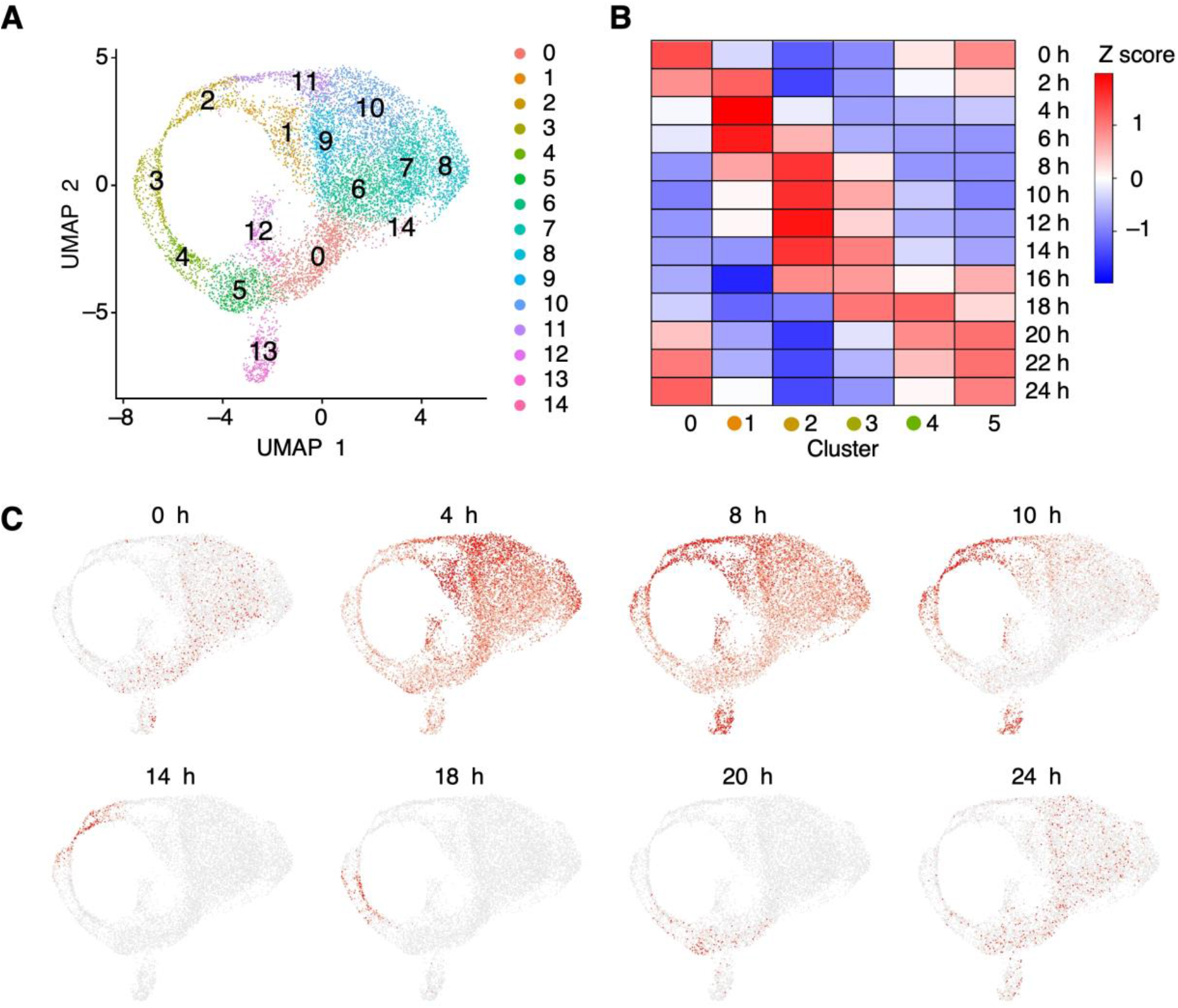
The Endogenous Diurnal Phase of Individual Cells Explains the Heterogeneity of Batch Cell Cultures. **(A)** UMAP plot for the 9,642 sequenced cells grown in Fe+ condition from experiment 1. The cells were separated into clusters by Seurat (Stuart et al., 2019) and are indicated by the color gradient, with the color key on the right side of the plot. **(B)** Heatmap representation of the average diurnal module scores associated with clusters 0-5 identified in (**A**). We calculated a diurnal module score for each cluster in 1 h phase bins based on diurnal phase data reported by Zones et al. (2015) of high confidence rhythmic genes, defined as the overlap of rhythmic genes from two recent studies (Zones et al., 2015; Strenkert et al., 2019). **(C)** UMAP plots of representative diurnal module scores for Fe+ cells from the first experiment.

Plotting phase module scores in UMAP plots also provided an opportunity to compare the phase distribution of Fe+ and Fe– cells. Indeed, although we collected cells at a single time-point, phase module scores reveal the endogenous phase of each cell, as a molecular timetable analysis would (Ueda et al., 2004). When we plotted diurnal module scores in UMAP plots for Fe– cells, we observed a similar pattern as that seen with Fe+ cells (Supplemental Figure 2). However, we discovered through a careful inspection of the UMAP plots that Fe– cells appeared to display a later diurnal phase relative to Fe+ cells, with more Fe– cells represented in the 8 h phase module plots, while Fe+ cells were more numerous in the 4 h module (Figure 3C, Supplemental Figure 2C). We interpret these results as suggestive of a delay in the circadian clock of the alga, reminiscent of the period lengthening effects observed under poor Fe nutrition in Arabidopsis (Hong et al., 2013; Chen et al., 2013; Salomé et al., 2013).

### Pseudo-time Construction Reveals the Phase Ordering of Batch Cultures

Until this point, we have considered one cell cluster as a unit and projected the diurnal module scores onto the clusters (Figure 3). To better characterize the rhythmic status of single cells, we selected clusters #0-#5 along the closed circle and used Monocle (Trapnell et al., 2014) to perform a pseudo-time analysis (Figure 4A). We ordered the cells into a continuous trajectory and assigned a pseudo-time point to each cell (Figure 4B, Supplementary Figure 3A, 3B). Next, we ordered cells by pseudo-time and plotted their associated diurnal module scores (Figure 4C). The pseudo-time trajectory started with cells from cluster #3, with a strong 14-18 h signature, that is shortly after cell division has occurred (Figure 3B, Supplemental Figure 3C). As pseudo-time increased, the trajectory progressed from cluster #3 through all other clusters in a counter-clockwise fashion, to end with cluster #2.

**Figure 4.**
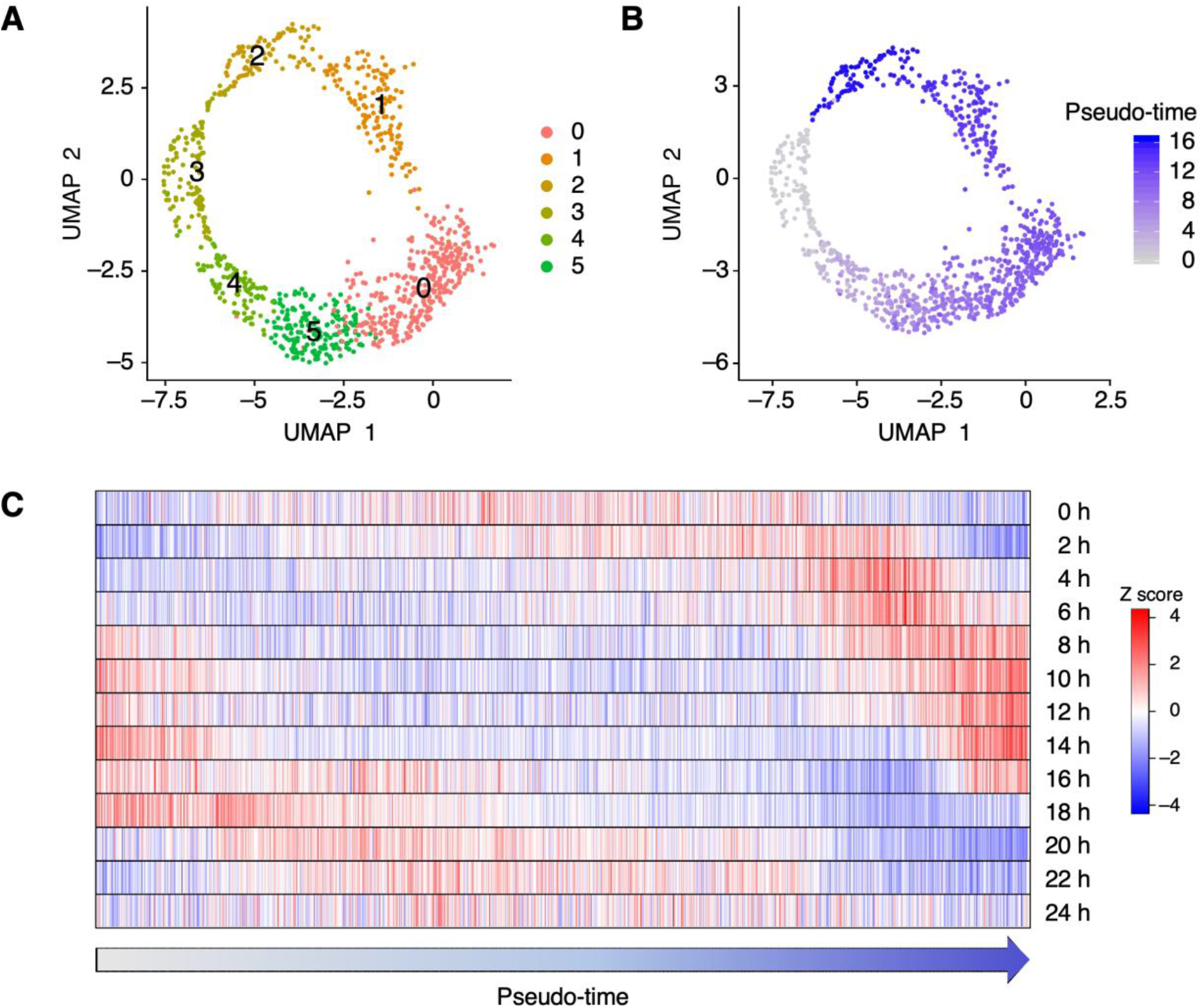
Pseudo-Time Construction Aligns Fe+ Cells Along the Diurnal Cycle. **(A)** UMAP plot of Fe+ cells from selected clusters progressing along the diurnal cycle shown in Figure 2A. Clusters are indicated by the color gradient, with the color key on the right side of the plot. **(B)** UMAP plot of the same Fe+ cell clusters, using pseudo-time as a graphical variable, indicated by the intensity of the blue dots. Note how pseudo-time starts at cluster #3 and runs counter clock-wise. **(C)** Heatmap representation of the diurnal module score in individual cells, ordered by pseudo-time. Each vertical bar corresponds to one individual cell.

That pseudo-time analysis tracked the diurnal phase bins underscores the essential contribution of rhythmic gene expression to the heterogeneity of Chlamydomonas cells in batch cultures.

### Effects of the Cell Wall on RNA Extractability and Quality for scRNA-seq

The protocols used for quantitative recovery of RNA from bulk Chlamydomonas cultures typically use ionic detergents and proteases, which are harsher than the typical extraction procedures used in the 10X pipeline. Therefore, we first used a *cw* mutant of Chlamydomonas for the previous analyses to facilitate RNA extraction and recovery. However, to apply these methods to natural field conditions or commercial pond situations, it would be useful to understand whether the same methodology might apply to walled cells. As a preliminary test, we incubated Chlamydomonas cells from strains with or without cell wall in the RNA extraction buffer used in the early steps before library construction. We also treated equal numbers of cells with 0.2% NP-40 and 2% SDS as positive controls for cell lysis, as seen by the release of chlorophyll from the cell pellet. As shown in Figure 5A, only the strain CC-5390, which lacks a cell wall, resulted in substantial lysis in the RT kit buffer, while we failed to observe signs of lysis with the other cell wall-containing strains CC-4532, CC4533 and CC-1690.

**Figure 5.**
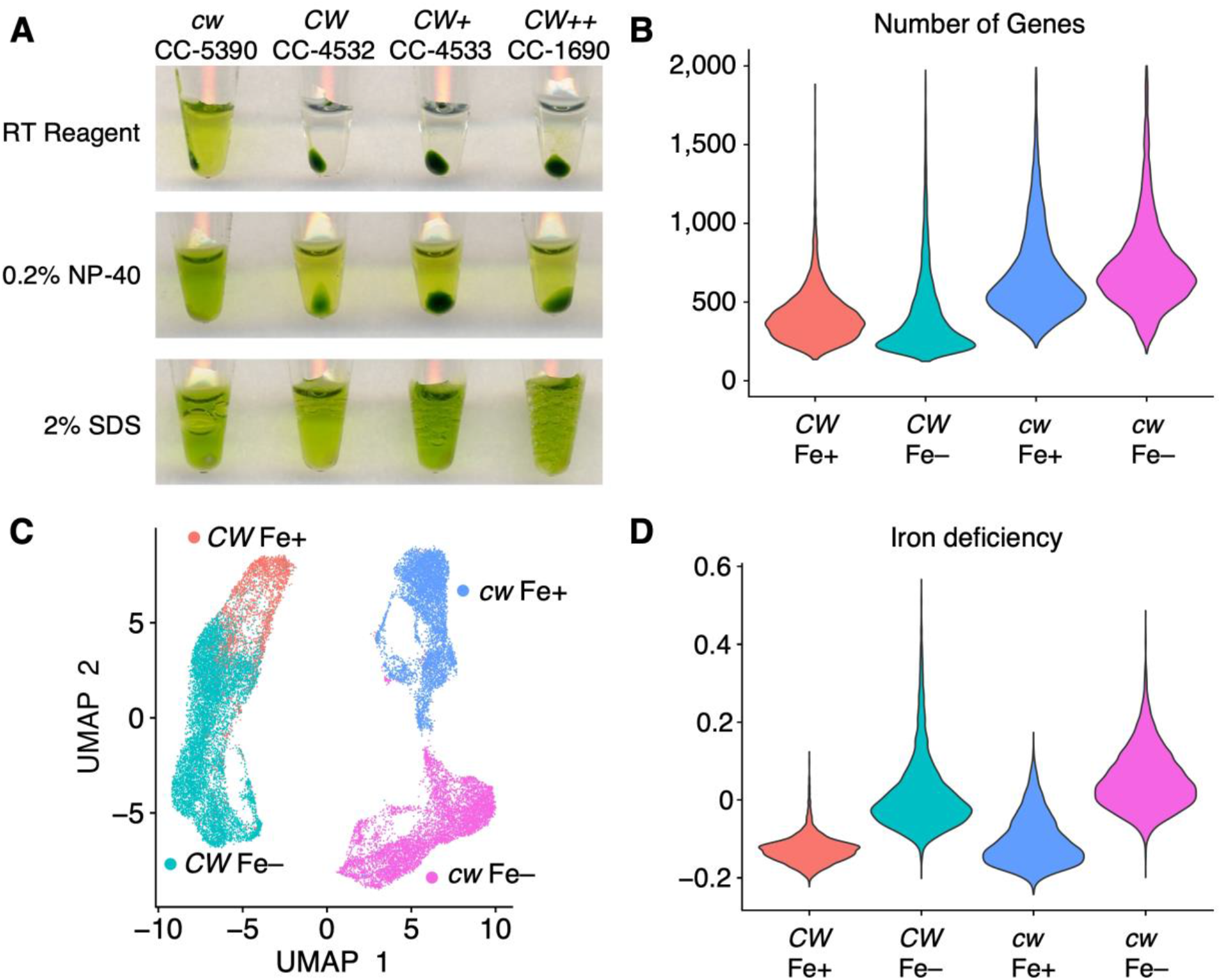
The Chlamydomonas Cell Wall Does not Block RNA Extraction for scRNA-seq Analysis. **(A)** Testing cell lysis with the RNA extraction buffer included in the 10X Chromium pipeline. We grew strains without (CC-5390, *cw*) or with (CC-4532, CC-4533, CC-1690, *CW*) a cell wall for 3 d in TAP medium before taking a 100 μL aliquot. After collection by centrifugation, cells were incubated with RT reagent (10X Genomics), 0.2% NP-40 or 2% SDS and incubated for 15 min before spinning cells again and taking the photograph. Strain CC-1690 has a thicker cell wall than CC-4532, as indicated by **(B)** Number of genes from which UMIs were detected in each sample. Strain CC-4532 was grown alongside strain CC-5390 during experiment 2 and treated in an identical manner. **(C)** UMAP plot of 24,795 sequenced cells from experiment 2, using Fe status and the presence of the cell wall as variables. **(D)** Iron deficiency module score associated with the cells shown in (**C**). For (**B-D**) Red: CC-4532 (*CW*) Fe+; teal: CC-4532 (*CW*) Fe–; blue: CC-5390 (*cw*) Fe+; magenta: CC-5390 (*cw*) Fe–.

Nevertheless, we selected strain CC-4532 (*CW*) for scRNA-seq on cells grown under iron-replete (Fe+) or Fe-starved (Fe–) conditions following the same methodology as for CC-5390. We processed both samples for GEMs production and library preparation. We successfully recovered sequenceable RNA from these samples and detected 2,814 Fe+ cells and 9,289 Fe– cells. When compared to CC-5390 (*cw*) strain grown under the same conditions, we collected data from fewer genes, reflecting some differences in RNA extractability or UMI formation in strains without (*cw*) or with (*CW*) a cell wall (Figure 5B).

To determine whether scRNA-seq captured the Fe nutritional status of strain CC-4532, we performed UMAP dimensionality reduction on CC-5390 (*cw*) and CC4532 (*CW*) samples grown side by side and treated in an identical manner as part of the second experiment. First, we noticed that the two strains clustered separately from each other, indicating strong transcriptomic differences correlated with the absence of the cell wall, strain-specific differences (Gallaher et al., 2015), or both (Figure 5C). Both strains formed distinct clusters corresponding to Fe+ and Fe– cells, demonstrating the applicability of scRNA-seq analysis to cell wall-containing algal strains, even without resorting to mechanical or enzymatic digestion. We did notice that the cluster formed by CC-4532 Fe– cells overlapped with that of CC-4532 Fe+ cells (Figure 5C). The Fe module score supported this observation (Figure 5D). We hypothesize that transferring cells from Fe-replete to Fe-starved conditions for 23 h was sufficient to induce a strong Fe deficiency response in CC-5390, whereas the cell wall-containing strain CC-4532 only partially depleted its Fe stores. Although this hypothesis has never been tested in two isogenic Chlamydomonas strains only differing at the *CW15* locus, empirical phenotyping of strains with and without cell walls under low Fe conditions is consistent with the higher sensitivity of cw strains to Fe deficiency (Allen et al., 2007; Gallaher et al., 2015).

## Discussion

We show that scRNA-seq can recapitulate bulk RNA-seq signatures and separate individual cells in non-overlapping clusters reflective of the growth condition they experienced (here, nutritional deficiency for Fe or N). In addition, we determine that Chlamydomonas cells grown in batch cultures retain substantial rhythmicity even after growing in constant light for weeks, contrary to common belief. This strong rhythmic component can explain much of the heterogeneity exhibited by individual Chlamydomonas cells in their transcriptional profile.

Using Arabidopsis and hairy bittercress (*Cardamine hirsuta*) as model systems, we had previously established that the Arabidopsis circadian clock responded to available Fe supply (Salomé et al., 2013). We and others showed that the circadian period lengthens under conditions of poor Fe nutrition, a phenotype that depended entirely on light-mediated chloroplast development (Salomé et al., 2013; Hong et al., 2013; Chen et al., 2013). One of several outstanding questions concerned the degree of evolutionary conservation of this response: do green single-cell algae such as Chlamydomonas adjust the period or phase of their circadian clock to the Fe status surrounding them? The comparison of diurnal phase module scores between Fe+ and Fe– Chlamydomonas cultures indicates that, in fact, Chlamydomonas cells do appear to adjust their diurnal phase as a function of their Fe status (Figure 3, Supplemental Figure 2). In addition, they do so in the same direction as Arabidopsis and hairy b ittercress, with a delay in diurnal phase under poor Fe nutrition conditions. Although our growth conditions did not specifically control for circadian behavior, these results nonetheless tentatively suggest that circadian Fe responses may be conserved between Chlamydomonas and Arabidopsis, opening new avenues for the systematic dissection of the underlying molecular mechanism by looking for conserved genes shared by the alga and the land plant.

Our Chlamydomonas cultures were maintained in constant light for weeks before sample collection. Yet, they showed a remarkable degree of synchronization that was not entirely expected. However, we independently reached the same conclusion from a deep re-analysis of hundreds of RNA-seq samples collected by our laboratory and the Chlamydomonas community over the past 10 years (Salomé and Merchant, submitted). Notably, one third of all bulk RNA-seq samples showed the same preferred diurnal phase as the single cell data described here. We hypothesize that Chlamydomonas cells may remain synchronized over such periods of time through two (non-mutually exclusive) hypotheses: 1) growing cells establish a population-wide phase, similar to quorum sensing in bacteria, that would maintain them in a synchronized state to share resources; 2) the manipulation of cells, for example the inoculation of the test cultures, acts as a synchronizing signal that persists for days. This latter possibility would be similar to a nutritional synchronization, such as serum shocks applied to mammalian cell cultures (Balsalobre et al., 1998). Cultures grown in flasks demand serial dilutions to remain in their exponential growth phase, making it difficult to determine the contribution of dilution to synchronization. By contrast, continuous flow bioreactors allow for absolute control of all parameters during cell culture, including cell density. We therefore envisage that the effect from inoculation as a resetting signal may be testable in bioreactors, thereby Chlamydomonas cells would be entrained by light-dark cycles and then released into constant light, all the while keeping the cell density low and constant. Samples may be collected every 12-24 h and processed for scRNA-seq, and the rhythmic components extracted as we did here, essentially following a molecular timetable approach applied to single cell populations (Ueda et al., 2004).

Our results also have commercial and ecological applications. Indeed, algal cells grown in large cultivation ponds may experience their surrounding environment differently as a function of pond depth, volume, cell density and turbulence. While bulk RNA-seq may help determine the average molecular and physiological phenotypes of cells collected at various depths and positions within the pond, the inherent variation between cells will be lost. By contrast, scRNA-seq offers a much more detailed picture of all cells within each sample, thus raising sensitivity by several orders of magnitude. Likewise, scRNA-seq applied to environmental samples collected in the wild may make it possible to describe algae in their native environment – what stresses they may experience and their interactions with other organisms with which they share the same ecological niche. Our results demonstrate that although cells lacking sufficient Fe or N stall along the cell cycle (Figure 2), they also express key stress marker genes that are inherently specific for each stress they may encounter. With carefully formulated gene lists and the calculation of the corresponding module scores, scRNA-seq may thus provide a unique opportunity to study Chlamydomonas (and other algae) in the wild. Chlamydomonas cells, just like yeast cells, can present a significant cell wall that might be considered a physical barrier for RNA extraction from single cells. In yeast, this technical limitation was resolved by adding the cell wall digesting enzyme zymolyase before (Jackson et al., 2020) or during (Jariani et al., 2020) the reverse transcription step of the same 10X Chromium Single Cell 30 v2 protocol we followed here. However, it should be noted that the authors did not attempt to generate scRNA-seq libraries from walled (undigested) yeast cells. Using chlorophyll release as a proxy for cell lysis, we similarly saw little lysis for the walled strain CC-4532; nevertheless, we detected hundreds of UMIs from this strain, indicating that Chlamydomonas strains of various cell wall thicknesses may be amenable to scRNA-seq. The Chlamydomonas cell wall is composed of a mixture of proteins and glycoproteins arranged in multiple layers, potentially limiting the use of cell wall digesting enzymes. A classic approach for the removal of the cell wall relies on autolysin, a zinc metalloprotease that is secreted by gametes during the initial stages of the algal sexual cycle. Treating cells with autolysin during the reverse transcription step may therefore mimic the initial stages of reproduction in Chlamydomonas, although this hypothesis can now be easily tested using the walled strain CC-4532 treated, or not, with autolysin. Another potential limitation to the use of autolysin is the difficulty associated with its purification from mating cells. A commercially-available protease would thus be preferable, such as alcalase, a commercial form of subtilisin that shows 35% identity with sporangin, the so-called hatching enzyme responsible for the digestion of the cell wall surrounding daughter cells before their release (Kubo et al., 2009; Hwang et al., 2019).

We only tested scRNA-seq on strains with no (like CC-5390) or moderately thick (like C-4532) cell wall. However, other laboratories focus on strains with a much more developed cell wall, for example CC-4533 (the wild-type background for a large insertional mutant library, Li et al., 2019) and CC-1690. The microfluidics pipeline from 10X Genomics now provides the perfect basis for a systematic comparison of RNA extraction efficiency across Chlamydomonas strains, with or without the addition of a protease during the reverse transcription step. The information gathered will also directly apply to wild isolates with walls, since the *cw* strains were all generated by mutagenesis in the laboratory (Hyams and Davies, 1972).

In conclusion, we showed that single-cell RNA-seq (scRNA-seq) can be applied to Chlamydomonas strains with or without a cell wall. In addition, scRNA-seq results recapitulated bulk RNA-seq data, indicating their reliability and the robustness of the Chlamydomonas transcriptome response to changes in its environment. Finally, we demonstrated that Chlamydomonas cells occupied a range of diurnal phases that may explain the heterogeneity exhibited by individual cells in bulk culture mode. By extracting diurnal data from single time-point scRNA-seq, we also observed a delay in the phase of the Chlamydomonas diurnal clock, suggesting that, just like land plants, algae may adjust the pace of their rhythms to Fe availability. The application of scRNA-seq to cultivation ponds and natural isolates will pave the way to a deeper understanding of the interactions between algae and their surroundings.

## Materials and Methods

### Growth Conditions

We used the *Chlamydomonas reinhardtii* strains CC-5390 (*cw15 arg7-8::ARG7 MT+*) and CC-4532 (*CW MT-*), which we procured from laboratory stocks. We grew all pre-cultures in Tris Acetate Phosphate (TAP) medium supplemented with micronutrients as described previously (Kropat et al., 2011), at 24°C in constant light (provided by a mixture of cool-white and warm-white fluorescent light bulbs, for a total Photon Flux Density ~50 μmol/m2/s) and under constant agitation (180 rpm) in an Innova-44R incubator.

In the first experiment, we started a pre-culture of strain CC-5390 in 50 mL TAP medium with 20 μM FeEDTA (iron replete conditions) at an initial cell density of 5 × 10^4^ cells/mL. After 5 d, we inoculated the test culture at the same initial cell density (5 × 10^4^ cells/mL), with 100 mL TAP medium + 20 μM FeEDTA in a 250 mL flask. After 5 d, we collected the cells by centrifugation for 3 min at 1,600*g* at room temperature using an Eppendorf centrifuge (model 5810 R), resuspended the pellet in 10 mL of fresh TAP medium (with 20 μM FeEDTA) and used 1 mL to inoculate a fresh flask containing 100 mL TAP medium + 20 μM FeEDTA, resulting in a 10-fold dilution of the culture. The next day, we pelleted the culture again across two 50 mL Falcon tubes, washed the pellets once with TAP medium without FeEDTA, and resuspended each pellet with either 50 mL TAP medium without FeEDTA (Fe– condition) or with 50 mL TAP medium + 20 μM FeEDTA (Fe+ condition) before transferring the cultures into fresh sterile 250 mL flasks and placing the flasks into the incubator. After 23 h of growth, we counted cell density in both cultures on a hemocytometer. Target cell density for scRNAseq analysis is 1,200 cells/μL: we therefore transferred 1.2 × 10^6^ cells/mL in a 1.5 mL Eppendorf tube, centrifuged the cells briefly on a tabletop centrifuge at 2,000rpm at room temperature. We resuspended the pellets into 1 × Phosphate Buffered Saline (PBS) with 0.04% BSA, placed the tubes on ice and covered them with aluminum foil. We walked to the Technology Center for Genomics and Bioinformatics at UCLA Pathology and Medicine (~5 min) for immediate processing, starting with Gel Bead in Emulsion (GEMs) formation.

For the second experiment, we started pre-cultures for CC-4532 and CC-5390 in 50 mL TAP medium + 20 μM FeEDTA at an initial cell density of 5 × 10^4^ cells/mL. After 3 d, we inoculated a new culture at the same initial cell density (4 flasks for CC-5390, 2 flasks for CC-4532). After another 3 d, we refreshed the cultures by 1:2 dilution with fresh TAP medium + 20 μM FeEDTA. The next day, we resuspended cultures in TAP without FeEDTA, TAP + 20 μM FeEDTA or TAP – nitrogen (CC-5390) or in TAP without FeEDTA or TAP + 20 μM FeEDTA (CC-4532), as described above. After 23 h of growth, we counted cells and proceeded as above.

### 10X Library Preparation, Sequencing, and Alignment

Cells were washed with PBS with 0.04% BSA, then counted with Countess II automated Cell Counter (Thermo Fisher). We loaded 10,000 cells onto the 10X Chromium Controller using Chromium Single Cell 3’ gene expression reagents (10X Genomics). The sequencing libraries were prepared following the manufacturer’s instructions (10X Genomics), with 12 cycles used for cDNA amplification and 12 cycles for library amplification. Library concentrations and quality were measured using Qubit ds DNA HS Assay kit (Life Technologies) and Agilent Tapestation 4200 (Agilent). The libraries were sequenced on a NextSeq500 platform as 2×50 paired-end reads to a depth of approximately 150 million reads per library (experiment 1), or using 2×50 paired-end reads, on an Illumina NovaSeq 6000 S2 platform to a depth of approximately 300 million reads per library (experiment 2). Raw reads were aligned to the Chlamydomonas genome (*Chlamydomonas reinhardtii* v5.5, Blaby et al., 2014) and cells were called using cellranger count (v3.0.2, 10X Genomics). Individual samples were aggregated to generate the merged digital expression matrix using the cellranger aggr pipeline (10X Genomics).

### Single-Cell RNA Sequencing Data Analysis

The R package Seurat (v3.1.2) (Stuart et al., 2019) was used to cluster the cells in the digital expression matrix. We filtered out cells with fewer than 100 genes or 300 unique molecular identifiers detected as low-quality cells. We divided the gene counts for each cell by the total gene counts for that cell, multiplied by a scale factor of 10,000, then natural-log transformed the counts. We used the FindVariableFeatures function from Seurat to select variable genes with default parameters. We used the ScaleData function from Seurat to scale and center the counts in the dataset. We performed Principal Component Analysis (PCA) on the variable genes, and selected 20 principle components for cell clustering (resolution = 0.5) and Uniform Manifold Approximation and Projection (UMAP) dimensionality reduction. We calculated module scores using the AddModuleScore function with default parameters.

### Calculation of the Diurnal Module Scores

We generated a list of diurnal signature genes by determining the overlap between rhythmic genes from two recent studies (Zones et al., 2015; Strenkert et al., 2019). The list contains 50 time points ranging from 0 to 24.5 h in ½ h interval. To calculate module scores from non-overlapping diurnal gene lists, we selected a 3 time-point interval that collapsed genes ½ h on either side of a given time-point. For example, the module score for diurnal phase 2 h was calculated using genes from the 1.5 h, 2 h and 2.5 h phase bins. Only the 0 h module score was calculated using genes from only two time points (0 h and 0.5 h). Dawn is taken as time 0 throughout.

### Pseudo-Time Trajectory Construction

We constructed pseudo-time trajectories using the R package Monocle (Trapnell et al., 2014). We extracted the raw counts for cells in the selected clusters and normalized them by the estimateSizeFactors and estimateDispersions functions with default parameters. We only retained genes with an average expression over 0.5 and detected in more than 10 cells for further analysis. We determined variable genes by the differentialGeneTest function with a model against the Seurat clusters. We determined the order of cells with the orderCells function, and constructed the trajectory with the reduceDimension function with default parameters.

### Compilation of Gene Lists for Module Score Analysis and scRNAseq Exploration

We assembled gene lists for the calculaton of module score by mining the literature. For the iron deficiency module score, we selected genes expressed > 10 FPKM and showing the stronger induction by Fe limitation from a comparison of RNA-seq data between Chlamydomonas CC-4532 grown in TAP medium + 0.25 μM FeEDTA and TAP medium + 20 μM FeEDTA (Urzica et al., 2012). We extracted the lipid biosynthesis and nitrogen gene lists from (Schmollinger et al., 2014). We ordered normalized expression data from a 48 h time-course in CC-4349 to identify genes that were induced in response to N deficiency (with normalized expression of 0 at 0 h and expression of 1 at 48 h), or repressed by N deficiency (or induced by N sufficiency, with normalized expression of 1 at 0 h and expression close to 0 at 48 h). The lists of lipid biosynthetic genes and ribosome protein genes were according to Supplemental Data Sets 14 and 9 from (Schmollinger et al., 2014), respectively The photosynthesis gene list include all nucleus-encoded genes from Supplemental Data Set 5 from (Strenkert et al., 2019). Cell cycle genes were obtained from Supplemental Data Set 4 of (Zones et al., 2015). Finally, we determined the diurnal phase of 10,294 high-confidence rhythmic genes by looking at the overlap between genes deemed to be rhythmic in two separate studies (Zones et al., 2015; Strenkert et al., 2019) and using the diurnal phase values from the 2015 work that had been recalculated for the 2019 study. Gene lists are provided as Supplemental Data Sets 1-10.

## SUPPLEMENTAL MATERIALS

**Supplemental Figure 1.** Modules scores for mitochondrial *RPGs* and lipid biosynthetic genes in cells from experiment 2. (Supports Figure 2).

**Supplemental Figure 2.** The Endogenous diurnal phase of individual cells explains the heterogeneity of batch cell cultures without iron. (Supports Figure 3).

**Supplemental Figure 3.** Pseudo-time construction aligns Fe+ cells along the diurnal cycle. (Supports Figure 4).

**Supplemental Table 1.** Summary of number of cells sequenced, number of genes and UMIs detected.

**Supplemental Data Set 1.** Fe deficiency module score gene list.

**Supplemental Data Set 2.** Nitrogen deficiency module score gene list.

**Supplemental Data Set 3**. Nitrogen sufficiency module score gene list.

**Supplemental Data Set 4.** Chloroplast ribosomal protein gene (*RPG*) module score gene list.

**Supplemental Data Set 5.** Cytosolic ribosomal protein gene (*RPG*) module score gene list.

**Supplemental Data Set 6.** Mitochondrial ribosomal protein gene (*RPG*) module score gene list.

**Supplemental Data Set 7.** Lipid biosynthesis module score gene list.

**Supplemental Data Set 8.** Cell division module score gene list.

**Supplemental Data Set 9.** Photosynthesis module score gene list.

**Supplemental Data Set 10.** Diurnal phase for high-confidence rhythmic genes.

## ACKNOWLEDGEMENTS

This work is supported by a cooperative agreement with the US Department of Energy Office of Science, Office of Biological and Environmental Research program under Award DE-FC02-02ER63421 (SM, MP), and in part (SM) by the Division of Chemical Sciences, Geosciences, and Biosciences, Office of Basic Energy Sciences of the U.S Department of Energy (DE-FD02-04ER15529). We thank Michael Mashock and other members of the Technology Center for Genomics & Bioinformatics (TCGB) at UCLA for preparing and sequencing 10X 3’ chromium single-cell libraries.

## AUTHOR CONTRIBUTIONS

PAS, MP and SSM designed the experiments. PAS grew all Chlamydomonas cultures and collected cells for scRNA-seq. FM mapped the reads to the Chlamydomonas genome and analyzed sequencing results. PAS provided gene lists to calculate module scores. PAS and FM wrote the manuscript with input from all authors.

## Data Availability

Sequence data from this article can be found at Phytozome under the following accession numbers:. *FEA1* (Cre12.g546550), *FEA2* (Cre12.g546600), *FRE1* (Cre04.g227400), *FOX1* (Cre09.g393150), *FTR1* (Cre03.g192050), *TEF22* (Cre12.g546500), *MSD3* (Cre16.g676150), *CDJ3* (Cre01.g009900), *CGLD27* (Cre05.g237050), *CTP1* (Cre16.g682369), *IRT1* (Cre12.g530400), *IRT2* (Cre12.g530350), *PHC1* (Cre17.g717900), *PHC21* (Cre02.g094450), *VSP1* (Cre11.g467710), *GAS28* (Cre11.g481600), *ACA4* (Cre10.g459200), *MTP1* (Cre03.g145087), *LCI6* (Cre12.g553350). Other genes used to calculate module scores are listed in Supplemental Data Sets 1-10.

scRNA-seq datasets were deposited at Gene Expression Omnibus at NCBI under the accession number GSE157580 (reviewers’ token: ytwdysegdpqthqr).

---

**Supplemental Figure 1. Modules scores for mitochondrial RPGs and lipid biosynthetic genes in cells from experiment 2.** (Supports Figure 2).

**(A)** Mitochondrial *RPG* module score for each sample.

**(B)** Lipid module score for each sample, using a gene list compiled by Schmollinger et al. (Schmollinger et al., 2014).

**Supplemental Figure 2. The Endogenous diurnal phase of individual cells explains the heterogeneity of batch cell cultures without iron.** (Supports Figure 3).

**(A)** UMAP plot of 9,642 sequenced cells from experiment 1 that were grown in Fe– condition for 23 h. The cells were separated into clusters by Seurat (Stuart et al., 2019) and are indicated by the color gradient, with the color key on the right side of the plot.

**(B)** Heatmap representation of the average diurnal module scores associated with clusters 0-8 identified in (**A**). We calculated a diurnal module score for each cluster in 1 h phase bins based on diurnal phase data reported by Zones et al. (2015) of high confidence rhythmic genes, defined as the overlap of rhythmic genes from two recent studies (Zones et al., 2015; Strenkert et al., 2019).

**(C)** UMAP plots of representative diurnal module scores for Fe– cells from the first experiment.

**Supplemental Figure 3. Pseudo-time construction aligns Fe+ cells along the diurnal cycle.** (Supports Figure 4).

**(A)** Trajectory plot of Fe+ cells from experiment 1, colored according to their constituent clusters. The five clusters shown here are the same clusters (#0-#5) shown in the UMAP plot in Figure 4.

**(B)** Trajectory plot of Fe+ cells from experiment 1, colored according to their pseudo-time. The plots in (**A**) and (**B**) are identical but colored based on the clusters they belong to (**A**) or according to their pseudo-time (**B**).

**(C)** Mean cell division scores for cells from clusters #0-#5 along pseudo-time coordinates. Note that a higher cell division module score is seen for clusters #3 and #2, consistent with their consecutive positions in UMAP plots.

## REFERENCES

Allen, M.D., Del Campo, J.A., Kropat, J., and Merchant, S.S. (2007). FEA1, FEA2, and FRE1, encoding two homologous secreted proteins and a candidate ferrireductase, are expressed coordinately with FOX1 and FTR1 in iron-deficient Chlamydomonas reinhardtii. Eukaryot. Cell 6: 1841–1852.

Bajhaiya, A.K., Dean, A.P., Zeef, L.A.H., Webster, R.E., and Pittman, J.K. (2016). PSR1 is a global transcriptional regulator of phosphorus deficiency responses and carbon storage metabolism in Chlamydomonas reinhardtii. Plant Physiol. 170: 1216–1234.

Balsalobre, A., Damiola, F., and Schibler, U. (1998). A serum shock induces circadian gene expression in mammalian tissue culture cells. Cell 93: 929–937.

Blaby-Haas, C.E., Castruita, M., Fitz-Gibbon, S.T., Kropat, J., and Merchant, S.S. (2016). Ni induces the CRR1-dependent regulon revealing overlap and distinction between hypoxia and Cu deficiency responses in: Chlamydomonas reinhardtii. Metallomics 8: 679–691.

Blaby-Haas, C.E. and Merchant, S.S. (2012). The ins and outs of algal metal transport. Biochim. Biophys. Acta – Mol. Cell Res. 1823: 1531–1552.

Blaby, I.K. et al. (2013). Systems-level analysis of nitrogen starvation-induced modifications of carbon metabolism in a Chlamydomonas reinhardtii starchless mutant. Plant Cell 25: 4305–4323.

Blaby, I.K. et al. (2014). The Chlamydomonas genome project: a decade on. Trends Plant Sci 19: 672–680.

Blaby, I.K., Blaby-Haas, C.E., Pérez-Pérez, M.E., Schmollinger, S., Fitz-Gibbon, S., Lemaire, S.D., and Merchant, S.S. (2015). Genome-wide analysis on Chlamydomonas reinhardtii reveals the impact of hydrogen peroxide on protein stress responses and overlap with other stress transcriptomes. Plant J. 84: 974–988.

Bogaert, K.A., Perez, E., Rumin, J., Giltay, A., Carone, M., Coosemans, N., Radoux, M., Eppe, G., Levine, R.D., Remacle, F., and Remacle, C. (2019). Metabolic, Physiological, and Transcriptomics Analysis of Batch Cultures of the Green Microalga Chlamydomonas Grown on Different Acetate Concentrations. Cells 8.

Chen, Y.Y., Wang, Y., Shin, L.J., Wu, J.F., Shanmugam, V., Tsednee, M., Lo, J.C., Chen, C.C., Wu, S.H., and Yeh, K.C. (2013). Iron is involved in the maintenance of circadian period length in Arabidopsis. Plant Physiol. 161: 1409–1420.

Eriksson, M., Moseley, J.L., Tottey, S., Del Campo, J.A., Quinn, J., Kim, Y., and Merchant, S. (2004). Genetic dissection of nutritional copper signaling in Chlamydomonas distinguishes regulatory and target genes. Genetics 168: 795–807.

La Fontaine, S., Quinn, J.M., Nakamoto, S.S., Dudley Page, M., Göhre, V., Moseley, J.L., Kropat, J., and Merchant, S. (2002). Copper-dependent iron assimilation pathway in the model photosynthetic eukaryote Chlamydomonas reinhardtii. Eukaryot. Cell 1: 736–757.

Gallaher, S.D., Fitz-Gibbon, S.T., Glaesener, A.G., Pellegrini, M., and Merchant, S.S. (2015). Chlamydomonas Genome Resource for Laboratory Strains Reveals a Mosaic of Sequence Variation, Identifies True Strain Histories, and Enables Strain-Specific Studies. Plant Cell 27: 2335–2352.

Gallaher, S.D., Fitz-Gibbon, S.T., Strenkert, D., Purvine, S.O., Pellegrini, M., and Merchant, S.S. (2018). High-throughput sequencing of the chloroplast and mitochondrion of Chlamydomonas reinhardtii to generate improved de novo assemblies, analyze expression patterns and transcript speciation, and evaluate diversity among laboratory strains and wild isolates. Plant J 93: 545–565.

Gasch, A.P. et al. (2017). Single-cell RNA sequencing reveals intrinsic and extrinsic regulatory heterogeneity in yeast responding to stress. PLoS Biol. 15.

González-Ballester, D., Casero, D., Cokus, S., Pellegrini, M., Merchant, S.S., and Grossman, A.R. (2010). RNA-Seq analysis of sulfur-deprived chlamydomonas cells reveals aspects of acclimation critical for cell survival. Plant Cell 22: 2058–2084.

Hong, S., Kim, S.A., Guerinot, M. Lou, and Robertson McClung, C. (2013). Reciprocal interaction of the circadian clock with the iron homeostasis network in arabidopsis. Plant Physiol. 161: 893–903.

Hwang, H.J., Kim, Y.T., Kang, N.S., and Han, J.W. (2019). A simple method for removal of the chlamydomonas reinhardtii cell wall using a commercially available subtilisin (Alcalase). J. Mol. Microbiol. Biotechnol. 28: 169–178.

Hyams, J. and Davies, D.R. (1972). The induction and characterisation of cell wall mutants of Chlamydomonas reinhardi. Mutat. Res. – Fundam. Mol. Mech. Mutagen. 14: 381–389.

Jackson, C.A., Castro, D.M., Saldi, G.A., Bonneau, R., and Gresham, D. (2020). Gene regulatory network reconstruction using single-cell rna sequencing of barcoded genotypes in diverse environments. Elife 9.

Jariani, A., Vermeersch, L., Cerulus, B., Perez-Samper, G., Voordeckers, K., Van Brussel, T., Thienpont, B., Lambrechts, D., and Verstrepen, K.J. (2020). A new protocol for single-cell RNA-seq reveals stochastic gene expression during lag phase in budding yeast. Elife 9: 1–22.

Kropat, J., Gallaher, S.D., Urzica, E.I., Nakamoto, S.S., Strenkert, D., Tottey, S., Mason, A.Z., and Merchant, S.S. (2015). Copper economy in Chlamydomonas: Prioritized allocation and eallocation of copper to respiration vs. Photosynthesis. Proc. Natl. Acad. Sci. U. S. A. 112: 2644–2651.

Kropat, J., Hong-Hermesdorf, A., Casero, D., Ent, P., Castruita, M., Pellegrini, M., Merchant, S.S., and Malasarn, D. (2011). A revised mineral nutrient supplement increases biomass and growth rate in Chlamydomonas reinhardtii. Plant J. 66: 770–780.

Kubo, T., Kaida, S., Abe, J., Saito, T., Fukuzawa, H., and Matsuda, Y. (2009). The chlamydomonas hatching enzyme, sporangin, is expressed in specific phases of the cell cycle and is localized to the flagella of daughter cells within the sporangial cell wall. Plant Cell Physiol. 50: 572–583.

Li, X. et al. (2019). A genome-wide algal mutant library and functional screen identifies genes required for eukaryotic photosynthesis. Nat. Genet. 51: 627–635.

Ma, X., Zhang, B., Miao, R., Deng, X., Duan, Y., Cheng, Y., Zhang, W., Shi, M., Huang, K., and Xia, X.Q. (2020). Transcriptomic and physiological responses to oxidative stress in a chlamydomonas reinhardtii glutathione peroxidase mutant. Genes (Basel). 11.

Malasarn, D., Kropat, J., Hsieh, S.I., Finazzi, G., Casero, D., Loo, J.A., Pellegrini, M., Wollman, F.A., and Merchant, S.S. (2013). Zinc deficiency impacts CO2 Assimilation and disrupts copper homeostasis in Chlamydomonas Reinhardtii. J. Biol. Chem. 288: 10672–10683.

Martin, N.C., Chiang, K.S., and Goodenough, U.W. (1976). Turnover of chloroplast and cytoplasmic ribosomes during gametogenesis in Chlamydomonas reinhardi. Dev. Biol.

Merchant, S.S. et al. (2007). The Chlamydomonas genome reveals the evolution of key animal and plant functions. Science (80-.). 318: 245–250.

Merchant, S.S., Allen, M.D., Kropat, J., Moseley, J.L., Long, J.C., Tottey, S., and Terauchi, A.M. (2006). Between a rock and a hard place: Trace element nutrition in Chlamydomonas. Biochim. Biophys. Acta – Mol. Cell Res. 1763: 578–594.

Moseley, J.L., Allinger, T., Herzog, S., Hoerth, P., Wehinger, E., Merchant, S., and Hippler, M. (2002). Adaptation to Fe-deficiency requires remodeling of the photosynthetic apparatus. EMBO J. 21: 6709–6720.

Park, J.J., Wang, H., Gargouri, M., Deshpande, R.R., Skepper, J.N., Holguin, F.O., Juergens, M.T., Shachar-Hill, Y., Hicks, L.M., and Gang, D.R. (2015). The response of Chlamydomonas reinhardtii to nitrogen deprivation: A systems biology analysis. Plant J. 81: 611–624.

Peltier, G. and Schmidt, G.W. (1991). Chlororespiration: An adaptation to nitrogen deficiency in Chlamydomonas reinhardtii. Proc. Natl. Acad. Sci. U. S. A. 88: 4791–4795.

Plumley, F.G. and Schmidt, G.W. (1989). Nitrogen-dependent regulation of photosynthetic gene expression. Proc. Natl. Acad. Sci. 86: 2678–2682.

Rodriguez, H., Haring, M.A., and Beck, C.F. (1999). Molecular characterization of two light-induced, gamete-specific genes from Chlamydomonas reinhardtii that encode hydroxyproline-rich proteins. Mol. Gen. Genet. 261: 267–274.

Salomé, P.A., Oliva, M., Weigel, D., and Krämer, U. (2013). Circadian clock adjustment to plant iron status depends on chloroplast and phytochrome function. EMBO J. 32.

Schmollinger, S. et al. (2014). Nitrogen-sparing mechanisms in Chlamydomonas affect the transcriptome, the proteome, and photosynthetic metabolism. Plant Cell 26: 1410–1435.

Shulse, C.N., Cole, B.J., Ciobanu, D., Lin, J., Yoshinaga, Y., Gouran, M., Turco, G.M., Zhu, Y., O’Malley, R.C., Brady, S.M., and Dickel, D.E. (2019). High-Throughput Single-Cell Transcriptome Profiling of Plant Cell Types. Cell Rep. 27: 2241–2247.e4.

Siersma, P.W. and Chiang, K.S. (1971). Conservation and degradation of cytoplasmic and chloroplast ribosomes in Chlamydomonas reinhardtii. J. Mol. Biol. 58.

Street, J.H. and Paytan, A. (2005). Iron, phytoplankton growth, and the carbon cycle. Met. Ions Biol. Syst. 43: 153–193.

Strenkert, D., Schmollinger, S., Gallaher, S.D., Salomé, P.A., Purvine, S.O., Nicora, C.D., Mettler-Altmann, T., Soubeyrand, E., Weber, A.P.M., Lipton, M.S., Basset, G.J., and Merchant, S.S. (2019). Multiomics resolution of molecular events during a day in the life of Chlamydomonas. Proc. Natl. Acad. Sci. U. S. A. 116: 2374–2383.

Stuart, T., Butler, A., Hoffman, P., Hafemeister, C., Papalexi, E., Mauck, W.M., Hao, Y., Stoeckius, M., Smibert, P., and Satija, R. (2019). Comprehensive Integration of Single-Cell Data. Cell 177: 1888–1902.e21.

Tilbrook, K., Dubois, M., Crocco, C.D., Yin, R., Chappuis, R., Allorent, G., Schmid-Siegert, E., Goldschmidt-Clermont, M., and Ulma, R. (2016). UV-B perception and acclimation in chlamydomonas reinhardtii. Plant Cell 28: 966–983.

Trapnell, C., Cacchiarelli, D., Grimsby, J., Pokharel, P., Li, S., Morse, M., Lennon, N.J., Livak, K.J., Mikkelsen, T.S., and Rinn, J.L. (2014). The dynamics and regulators of cell fate decisions are revealed by pseudotemporal ordering of single cells. Nat. Biotechnol. 32: 381–386.

Ueda, H.R., Chen, W., Minami, Y., Honma, S., Honma, K., Iino, M., and Hashimoto, S. (2004). Molecular-timetable methods for detection of body time and rhythm disorders from single-time-point genome-wide expression profiles. Proc. Natl. Acad. Sci. 101: 11227–11232.

Urzica, E.I., Casero, D., Yamasaki, H., Hsieh, S.I., Adler, L.N., Karpowicz, S.J., Blaby-Haas, C.E., Clarke, S.G., Loo, J.A., Pellegrini, M., and Merchant, S.S. (2012). Systems and trans-system level analysis identifies conserved iron deficiency responses in the plant lineage. Plant Cell 24: 3921–3948.

Waffenschmidt, S., Woessner, J.P., Beer, K., and Goodenough, U.W. (1993). Isodityrosine cross-linking mediates insolubilization of cell walls in chlamydomonas. Plant Cell 5: 809–820.

Wittkopp, T.M. et al. (2017). Bilin-dependent photoacclimation in Chlamydomonas reinhardtii. Plant Cell 29: 2711–2726.

Zhang, T.Q., Xu, Z.G., Shang, G.D., and Wang, J.W. (2019). A Single-Cell RNA Sequencing Profiles the Developmental Landscape of Arabidopsis Root. Mol. Plant 12: 648–660.

Zhang Y., Wang J., Sang Y., Jin S., Wang X., Kumar Azad G., McCormick M.A., Kennedy B.K., Li Q., Wang J., Z.X. and H.Y. (2020). Single-cell RNA-seq reveals early heterogeneity during ageing in yeast. Biorxiv.

Zones, J.M., Blaby, I.K., Merchant, S.S., and Umen, J.G. (2015). High-Resolution Profiling of a Synchronized Diurnal Transcriptome from Chlamydomonas reinhardtii Reveals Continuous Cell and Metabolic Differentiation. Plant Cell 27: 2743–2769.

