## Supplemental Materials for "Single-Cell RNA Sequencing of Batch Chlamydomonas Cultures Reveals Heterogeneity in their Diurnal Cycle Phase"

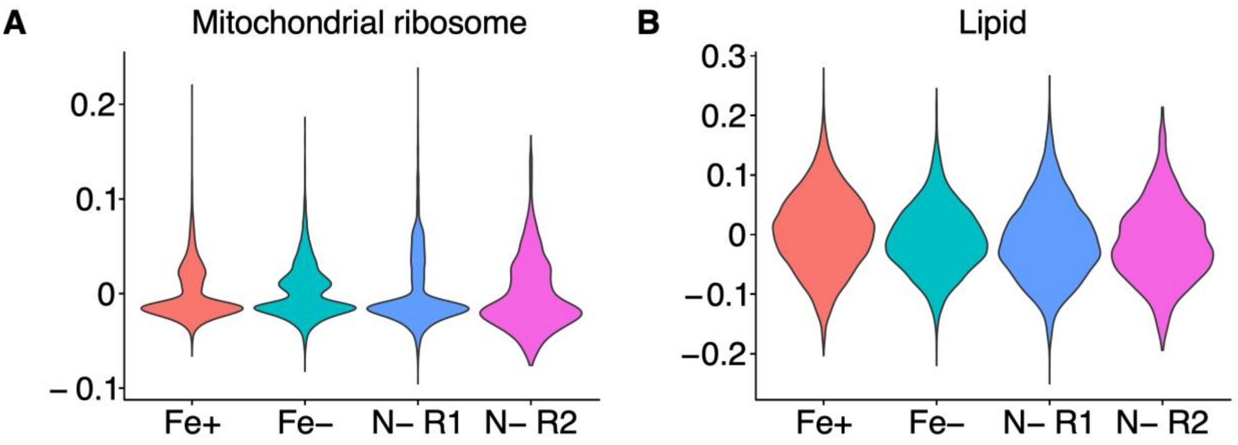


**Supplemental Figure 1. Modules scores for mitochondrial RPGs and lipid biosynthetic genes in cells from experiment 2.** (Supports Figure 2).

**(A)** Mitochondrial *RPG* module score for each sample.

**(B)** Lipid module score for each sample, using a gene list compiled by Schmollinger et al. (Schmollinger et al., 2014).

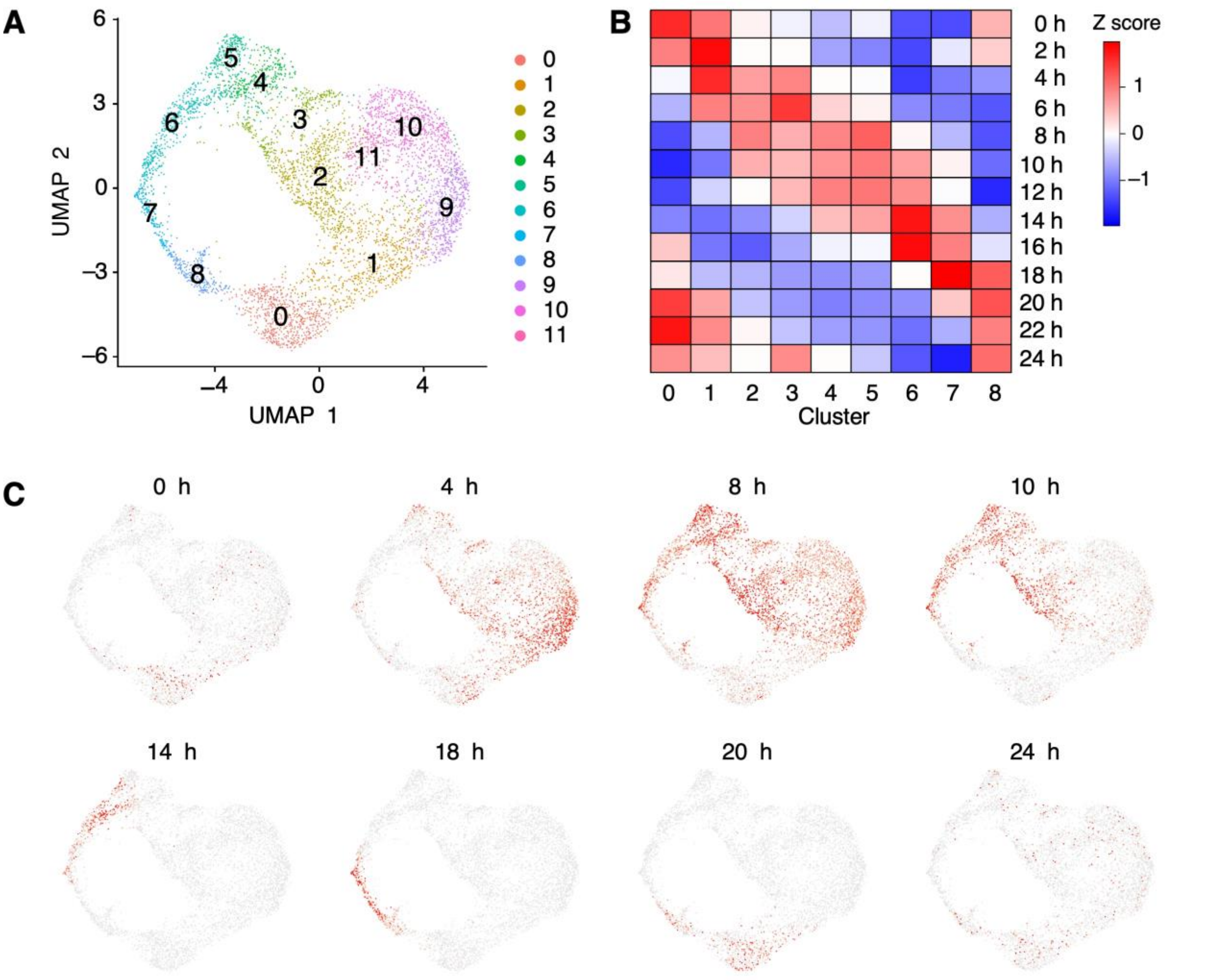


**Supplemental Figure 2. The Endogenous diurnal phase of individual cells explains the heterogeneity of batch cell cultures without iron.** (Supports Figure 3).

**(A)** UMAP plot of 9,642 sequenced cells from experiment 1 that were grown in Fe– condition for 23 h. The cells were separated into clusters by Seurat (Stuart et al., 2019) and are indicated by the color gradient, with the color key on the right side of the plot.

**(B)** Heatmap representation of the average diurnal module scores associated with clusters 0-8 identified in (**A**). We calculated a diurnal module score for each cluster in 1 h phase bins based on diurnal phase data reported by Zones et al. (2015) of high confidence rhythmic genes, defined as the overlap of rhythmic genes from two recent studies (Zones et al., 2015; Strenkert et al., 2019).

**(C)** UMAP plots of representive diurnal module scores for Fe– cells from the first experiment.

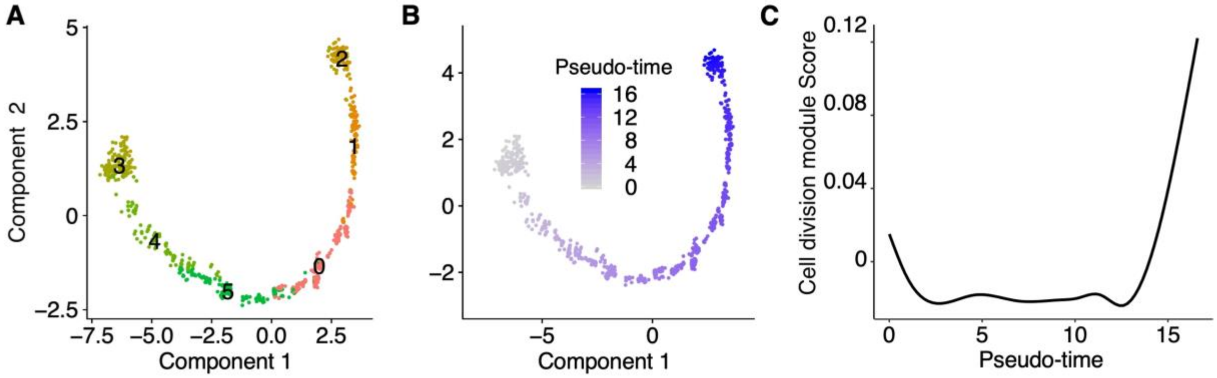


**Supplemental Figure 3. Pseudo-time construction aligns Fe+ cells along the diurnal cycle.** (Supports Figure 4).
**(A)** Trajectory plot of Fe+ cells from experiment 1, colored according to their constituent clusters. The five clusters shown here are the same clusters (#0-#5) shown in the UMAP plot in Figure 4.
**(B)** Trajectory plot of Fe+ cells from experiment 1, colored according to their pseudo-time. The plots in (**A**) and (**B**) are identical but colored based on the clusters they belong to (**A**) or according to their pseudo-time (**B**).
**(C)** Mean cell division scores for cells from clusters #0-#5 along pseudo-time coordinates. Note the higher cell division module score for clusters #3 and #2, consistent with their consecutive positions in UMAP plots.

**Supplemental Table 1.** Summary of number of cells sequenced, number of genes and UMIs detected.

| Experiment | Sample | Strain | Number cells | UMIs per cell | Genes per cell |
|---|---|---|---|---|---|
| 1 | Fe+ | CC-5390 (*cw*) | 9,517 | 3,009 | 827 |
| 1 | Fe– | CC-5390 (*cw*) | 9,748 | 3,716 | 828 |
| 1 | Mix | CC-5390 (*cw*) | 9,425 | 3,297 | 815 |
| 2 | Fe+ | CC-5390 (*cw*) | 7,960 | 4,044 | 630 |
| 2 | Fe– | CC-5390 (*cw*) | 6,656 | 5,380 | 747 |
| 2 | Fe+ | CC-4532 (*CW*) | 2,814 | 3,961 | 409 |
| 2 | Fe– | CC-4532 (*CW*) | 9,289 | 2,757 | 414 |
| 2 | N– R1 | CC-5390 (*cw*) | 3,028 | 2,034 | 578 |
| 2 | N– R2 | CC-5390 (*cw*) | 1,496 | 3,924 | 1,035 |
| | | Total | 59,933 | | |
